# CRISPR Knock-in Designer: automatic oligonucleotide design software to introduce point mutations using CRISPR/Cas9

**DOI:** 10.1101/2021.04.11.439384

**Authors:** Sergey V. Prykhozhij, Vinothkumar Rajan, Kevin Ban, Jason N. Berman

**Affiliations:** Children’s Hospital of Eastern Ontario Research Institute, Ottawa, ON, Canada; Biological Sciences Platform, Sunnybrook Research Institute, Toronto, ON, Canada; Departments of Pediatrics and Cellular and Molecular Medicine, University of Ottawa, Ottawa, ON, Canada

## Abstract

Knock-in of precise point mutations into protein-coding genes has been one of the earliest and most important applications of Clustered Regularly Interspaced Palindromic Repeats (CRISPR)/Cas9. The ability to perform such precise gene editing is crucial to interrogate the function of specific protein residues and to create models of human diseases caused by protein amino acid changes. The homologous protein residues can be mutated in model animal species, and the consequences of these mutations can be studied, leading to a better understanding of the disease in question. Design of point mutation knock-in strategies has been a combination of manual steps assisted by several computational tools resulting in a time-consuming process and preventing a single rapid and integrated solution. We have therefore designed CRISPR Knock-in Designer, which can perform rapid and automatic design of point mutation knock-in DNA oligonucleotides upon provision of the mutation, a guide RNA, and essential identifier or sequence information. The tool supports most experimentally established CRISPR types and has multiple options for the resulting oligonucleotides to satisfy the needs of most users. We also provide allele-specific PCR-based and restriction enzyme-based genotyping strategies as part of the program output. CRISPR Knock-in Designer adjusts to the genomic context of any target codon and tries to design knock-in strategies when a codon straddles two exons, a situation we explored in whole genomes of several model species. CRISPR Knock-in Designer output can also be adapted for use with some of the newer Prime Editing design tools to facilitate the introduction of a specific mutation sequence using this advanced technology.

## Introduction

Adaptation of the Clustered Regularly Interspaced Palindromic Repeats (CRISPR)/Cas9 bacterial system to perform genome editing represents a great triumph of molecular biology (1–5). Briefly, this system is based on the ability of Cas9 or other Cas nucleases to cut specific regions of double-stranded DNA (dsDNA) with the specificity provided by 18-20bp single guide RNAs (sgRNA) or other CRISPR RNAs (crRNA) that can be designed. Guide RNA (gRNA) consists of a spacer region and a Protospacer Adjacent Motif (PAM), together defining where the cut site will be located. Once the target DNA site is cut, the cell typically uses its DNA repair mechanisms to bring together the cut DNA ends using the error-prone nonhomologous end-joining (NHEJ) pathway to create insertions/deletions (indels) (6). By contrast, researchers provide template DNA molecules that can divert DNA repair toward precise homology-directed repair (HDR) pathways to introduce specific modifications.

Generation of precise point mutations for basic research and disease modeling studies is an important contribution of the burgeoning CRISPR/Cas9 technology, where Cas9-sgRNA ribonucleoprotein introduces the cut into the complementary genomic DNA and a donor template can then be used for introducing the desired mutant sequence. Single-stranded oligodeoxynucleotides (ssODNs), or oligos for short, are currently the donor templates of choice for point mutation knock-ins. HDR includes the following subpathways: Double-strand Break Repair (DSBR), Synthesis-dependent Strand Annealing (SDSA) and single-strand DNA Incorporation (ssDI) (7). Due to being single-stranded, ssODNs cannot utilize DSBR, and the balance between SDSA and ssDI was investigated by a point mutation knock-in study in human cells (7). SDSA is the dominant knock-in pathway when a double-strand break is introduced, whereas the ssDI pathway, involving direct incorporation of the oligonucleotide fragment, was mainly active in a PAM-in (PAMs toward each other) paired nickase design generating protruding 3’ ends. CRISPR and ssODN-based point mutation knock-ins have been performed in human cells (8), mice (9), pigs (10), zebrafish (11, 12), *Drosophila melanogaster* (13), *Caenorhabditis elegans* (14), and likely most other species where CRISPR/Cas9 is applicable.

The main challenges for point mutation knock-ins remain their low efficiency relative to a high percentage of insertions and deletions (indels). We have optimized this technology in zebrafish to generate cancer-relevant point mutants in *tp53* and several other genes (12) and have reviewed other studies which applied very similar techniques to introduce specific knock-in mutations (15). The primary ways to improve knock-in using oligos is to ensure the minimum distance between the cut site and the target mutation, use correct asymmetric designs from both DNA strands, incorporate phosphorothioate nucleotides into the oligo ends and ensure that the efficiency of Cas9-sgRNA reagents is optimal (16). Such optimization usually leads to good results, but the overall average efficiency remains below 10% in the context of zebrafish and several other systems. zLOST (zebrafish long single-stranded DNA template) method for CRISPR/Cas9-based point mutation knock-ins with enzymatically-synthesized long single-stranded DNA of 300-500 nucleotides as templates has recently been shown to have higher efficiencies in zebrafish compared to shorter oligonucleotides or double-stranded DNA molecules (17). This result is similar to the Easi-CRISPR method in mice, which has been mainly used for fluorescent protein tag insertions (18).

One component missing in the process of point mutation knock-in design is a computational pipeline for designing oligos for any desired point mutation. Thus far, our go-to approach for knock-in design has been to manually design the oligos with the aid of some standard text processing and additional bioinformatics tools. However, this approach is time-consuming and may not result in optimal results. This realization led us to implement the computational design pipeline for point mutations in protein-coding genes. This pipeline, further referred to as *CRISPR Knock-in Designer*, works with gene sequence data provided directly or retrieves these data based on a transcript identifier. It can introduce the desired point mutations, optionally mutate the PAM or guide RNA spacer sequence, and introduce further silent mutations for inserting unique restriction sites suitable for genotyping. Any other unique restriction sites introduced by mutations are also identified and presented in the output. The program also designs allele-specific PCR (AS-PCR) assays from both sides of the target modification.

Although most codons lie within exons, some codons straddle two exons leading to the notion of exon and intron phases. For introns, phase indicates if an intron does not split the next codon (phase 0) or occurs after the first or the second nucleotide of the next codon (phases 1 or 2, respectively). Exon phases are defined by the phases of flanking introns. Comparison of intron positions in homologous genes across large evolutionary distances has broadly supported the early origin of introns during eukaryotic evolution (19). CRISPR Knock-in Designer takes care to recognize the position of any target codon with respect to the exons and tests whether the input mutation is possible. This mutability test for all possible codon-amino acid residue combinations resulted in mutability matrices. We have further explored the prevalence of such two-exon straddling codons in protein-coding genes of several animal model species and found their higher prevalence in chordate species than in *D. melanogaster* and *C. elegans*.

Prime Editing (PE) is a new genome editing technology relying on a fusion of a Cas9 nickase with a reverse transcriptase enzyme to perform edits based on the information encoded in a prime editor guide RNA (pegRNA) (20). In the PE2 strategy, Cas9 nickase with pegRNA binds to the complementary strand and cuts the opposite strand, the pegRNA primer-binding site binds to the released single-stranded DNA, and the reverse transcriptase enzyme copies the edited sequence from the pegRNA into the single strand of DNA, which is then repaired with the edit being stably introduced into the DNA by NHEJ. In the PE3b strategy, the nickase with the regular guide RNA cuts the unedited strand thus promoting repair using the edited DNA strand (20). PE could replace ssODN-based CRISPR knock-ins where it is applicable and advantageous. Nevertheless, the current CRISPR knock-in techniques remain viable and useful for the foreseeable future, and the resulting experience may facilitate PE design and adoption. Multiple prime editing design tools (pegFinder (http://pegfinder.sidichenlab.org/), PrimeDesign (https://drugthatgene.pinellolab.partners.org/)) require that the desired mutations be pre-designed or labeled in their genomic context. To make CRISPR Knock-in Designer output compatible with such tools, we provide genomic sequences of up to 500 bp with the designed mutations already incorporated as output files. In summary, CRISPR Knock-in Designer is a novel software to design CRISPR point mutation knock-in strategies and can be employed for prime editing design applications.

### Implementation

#### Data input overview

The process of introducing point mutations using CRISPR/Cas9 into the genomic DNA of any model species or cell line consists of multiple steps. First, an active guide RNA needs to be identified close to the target site. This is an experimental procedure familiar to all the users of the CRISPR/Cas9 technology and is required for the design of oligonucleotides to introduce point mutations. We have focused on missense or nonsense mutations in the protein-coding genes because other precise point mutations may be too specific for a general design tool and imprecise indel mutations induced by CRISPR/Cas9 strategies typically do not require any oligonucleotides for their generation. CRISPR Knock-in Designer automates multiple steps needed to design donor oligonucleotides. The users need to provide multiple inputs (Fig. 1). The target mutation needs to be defined with a gene name, and a mutation string such as X123Y, where X is a single-letter amino acid (AA) code and Y can be an AA code or stop codon (*). Sequence input to the program can be provided manually or retrieved automatically using the provided Ensembl Transcript ID. The manual input requires more effort since the user must provide up to 4 relevant sequences. Coding DNA sequence (CDS) containing all codons of a protein and the target exon sequence are essential for the process since CDS is used to verify the target codon as well as the exon, where the target codon is located. Optionally, the user may also need to provide 5’ and 3’ flanking sequences, which may be introns or other genomic sequences. In contrast to the manual data entry, the automatic data entry only requires an Ensembl Transcript identifier (ID) to retrieve relevant sequences. Ensembl REST API is then used to infer the species and retrieve all relevant sequences. This allows the tool to function universally in species whose genomes are available in Ensembl, and for still other species, the design process can be run in the manual data entry mode.

**Figure 1.**
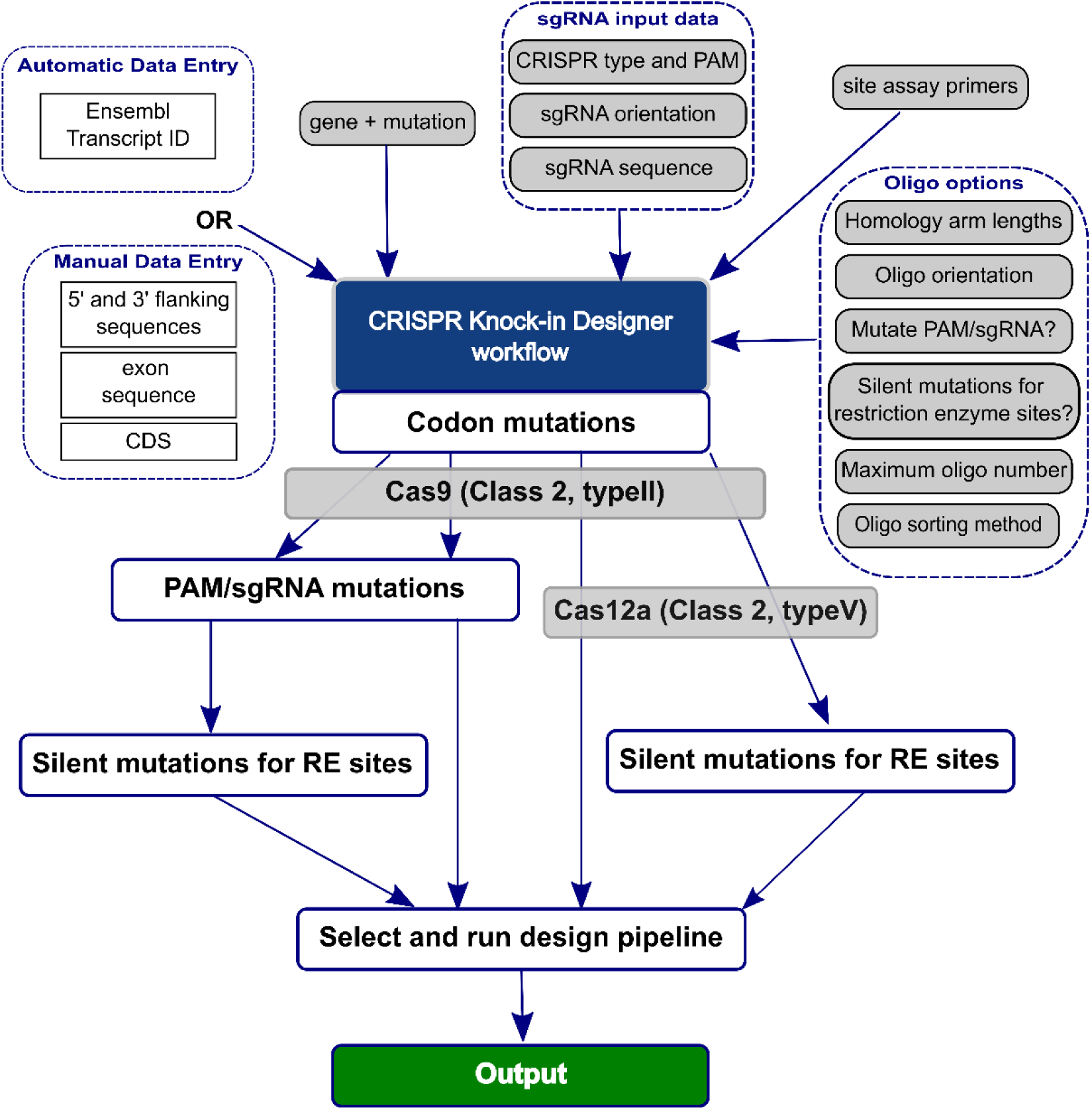
Overview of the CRISPR Knock-in Designer algorithm. The protein-coding gene mutation is defined here by the gene symbol and a mutation string such as X123Y (gene + mutation) in single-letter amino acid code. The relevant sequences are either retrieved automatically using the Ensembl Transcript ID input (Automatic Data Entry) or provided manually. The Manual Data Entry option requires coding DNA sequence (CDS) and exon sequence of the target residue codon, the flanking sequences being optional depending on the primer binding sites. Another important input is site assay (amplicon around the target site) primers that define the genomic region for the genotyping assay designs. Single guide RNA (sgRNA) input data consist of the CRISPR type and PAM parameter, sgRNA orientation and sequence. The Oligo options include homology arm length, oligo orientation, “Mutate PAM/sgRNA?” (whether to mutate the PAM or sgRNA sequence), “Silent mutations for restriction enzyme sites?” (whether to introduce restriction sites by synonymous mutations near the target codon), Maximum oligo number and Oligo sorting method. The program then performs the basic target codon mutation procedure followed by additional mutation procedures that are chosen based on the oligo options and the CRISPR type. All paths of program execution are shown and those that can be chosen based on the CRISPR type are overlaid with the CRISPR type label. However, in each run, only a single design pipeline can be chosen, which results in the output of the designed oligonucleotides and the corresponding genotyping strategies.

Further, the user must provide “site assay primers” (Fig. 1) denoted this way to reflect their use to verify the activity of gRNAs at their binding sites. Since exons are frequently short (about 100 bp) but can also be much larger, and the target codons can be located anywhere inside them, the primers used to verify the activity of guide RNAs can bind either the exon or the flanking sequences with four possible binding site combinations and the corresponding requirements for input sequences during the manual data entry (Fig. 2A). The automatic sequence retrieval and the corresponding design procedures handle this issue automatically by always retrieving the exon and 600-bp flanking sequences on each side followed by extracting the site assay amplicon sequence.

**Figure 2.**
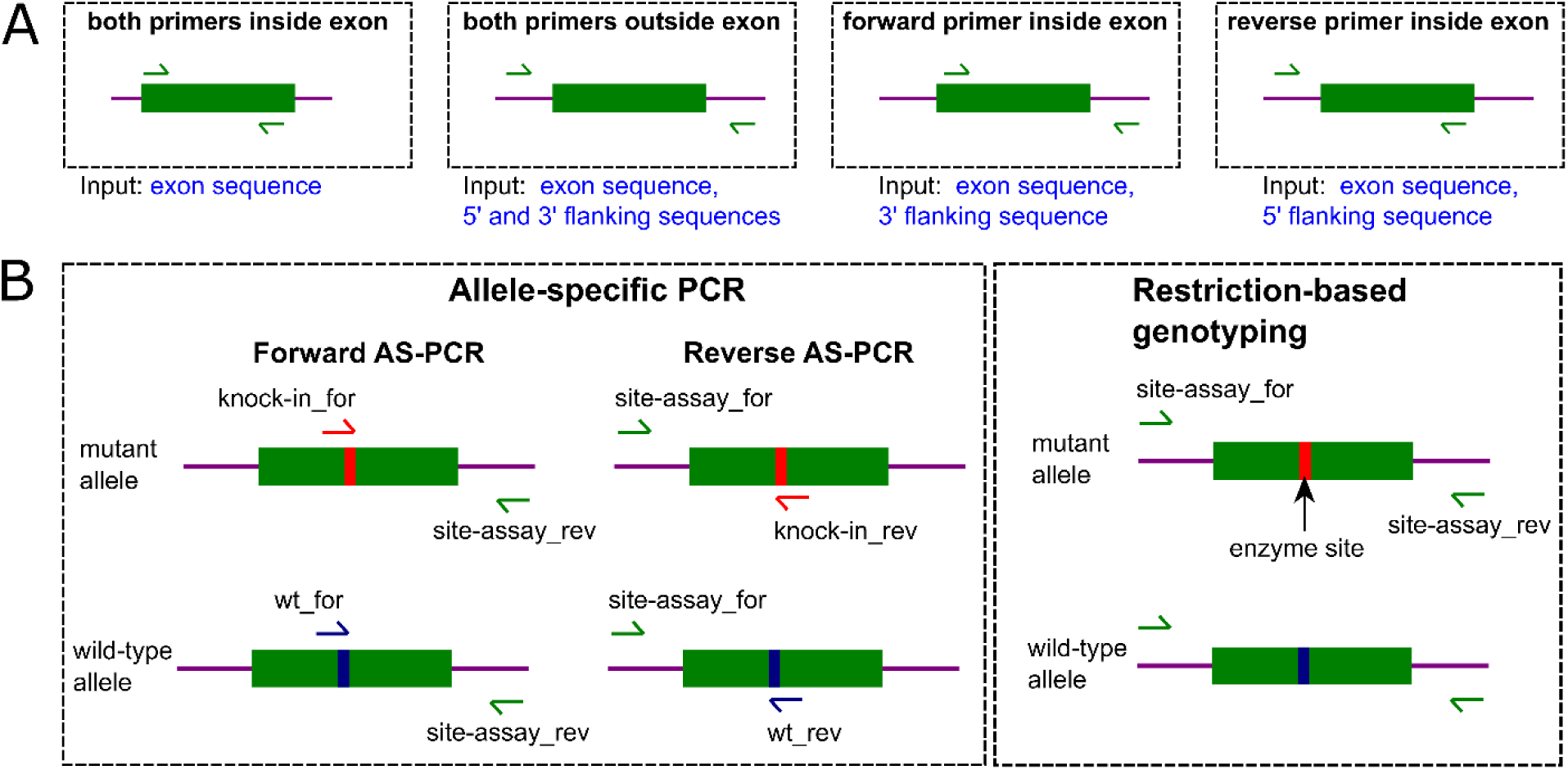
Site assay primer binding sites, sequence input requirements and genotyping strategies. **A**. Locations of the forward and the reverse site assay primers provided by the user are visualized here with respect to the exon (green block) and the flanking sequences (magenta lines). These locations determine the required manual data entry inputs as indicated by the text under the dashed boxes resulting in four possible input sequence combinations. **B**. Mutations introduced by point mutation knock-ins can be genotyped in two ways: by allele-specific PCR (AS-PCR) or using restriction enzyme-based genotyping. In the AS-PCR approach, one can design either forward (forward AS-PCR) or reverse (reverse AS-PCR) primers matching the mutant or wild-type alleles and then run these assays to detect the introduced point mutations and to verify them against the wild-type allele. After the point mutation knock-in, one may also introduce a specific restriction site and then use the corresponding enzyme to genotype the modified cells or organisms. This approach is best applied when modification efficiency is high or after the modification has been completed and the purpose is to distinguish various genotypes after Mendelian gene distribution following crosses.

CRISPR Knock-in Designer assumes the availability of an active gRNA to design an oligonucleotide for a point mutation knock-in, so it is required that the user provides the sequence, its orientation, and the PAM sequence defining the nuclease enzyme. We focused on the class 2 type II enzymes such as *Streptococcus pyogenes* Cas9 and its engineered mutants with altered PAM sequences, *Staphylococcus aureus* Cas9 and its KKH mutant as well as class 2 type V Cas12 enzymes. For the purposes of point mutation knock-ins, differences in PAMs of Cas9 and Cas12a enzymes are likely to make them complementary since they will have their respective applicability in more GC and AT-rich genomic regions. Another important consideration is the difference in the cut sites between the nucleases since Cas12a cuts at approximately 18 nt from the PAM on one strand and at 23 nt on the other strand leaving 5’ overhangs, whereas Cas9 enzymes make blunt cuts 3 nt away from the PAM. Therefore, Cas12a PAMs would typically have to be more distant from the target codons than would be the case for Cas9 if both are to achieve high knock-in efficiency.

The oligo options input section allows the user to select oligo homology arm lengths and the orientation of the oligos. Further, the user can select whether to perform synonymous codon mutations to inactivate PAM or gRNA site binding. This will prevent re-cutting after the sequence is copied from the oligo to the genomic DNA. This option is mainly applicable to Cas9 designs since oligonucleotide-based knock-in efficiency is known to drop dramatically with the distance from the cut site (9) which led us not to apply this option to Cas12a designs. We also provide an option to introduce restriction sites with synonymous mutations of codons located next to the target codon (“Silent mutations for restriction enzyme sites?”). This option will strongly facilitate genotyping of the resulting point mutations. Given that for some strategies the number of possible designed oligonucleotides can be large (>20), we provided the “Maximum oligo number” option as a slider input. The final option is “Choose how to sort oligos”, which allows the user to do no sorting, random sorting or sorting by the average of absolute distances between the target codon mutation and additional mutations. The random sorting option is mainly useful to mix oligos with different replacement codons. The “average mutation-codon distance” option sorts oligos in such a way that oligos with more mutation clustering will occupy top positions in the list thus increasing the likelihood that all the designed mutations will be introduced into the genome together and improving the specificity of their detection.

### Input data validation

After the user provides the input, the program needs to perform input data validation. This prevents unhandled errors in the program execution or erroneous results if the program still manages to proceed with partially incorrect data. shinyFeedback provides the first line of data checking by verifying that the input is not empty, that the input sequences are DNA sequences, that they conform to the expected pattern, satisfy minimum length requirements or, in the case of coding sequences that their lengths are divisible by 3. For the mutation input, we validate that the mutation input is not empty and conforms to the pattern “XN{1,5}Y”, where X is a single-letter amino acid code in the upper or lower case, N{1,5} is an amino acid position from 1 to 99999 and Y is either a single-letter amino acid code or a stop codon (*).The retrieval of sequence data from Ensembl REST API is initially assumed to be correct but is validated at the end of the process when the mutation target codon is mapped to the retrieved exons to determine if this codon is fully within an exon sequence or straddles two exons. If this mapping fails, the validation error is raised, and the program terminates. For the manual input, all sequence input is validated to consist of DNA characters, the CDS and the exon sequence are ensured not to be empty, and CDS length must be divisible by 3. After the sequence input is completed by either automatic retrieval or manual input, the target codon is validated to encode the amino acid indicated in the mutation string. This is important because the corresponding AA residues may have different positions in different transcript isoforms, and the program needs to ensure the correct mapping between the CDS and the target codon position. The exon sequence is also mapped by sequence alignment to CDS and successful alignment is validated. Primer inputs are likewise ensured not to be empty, and their validation is performed by successful alignment to the genomic sequence around the mutation. Mapping of the input sgRNA sequence to the input genomic sequence by alignment is determined by both the sequence and orientation of sgRNA, so these inputs are validated jointly to avoid ambiguity. Finally, the type of CRISPR is validated by retrieving the PAM sequence from the genomic sequence and matching it to the expected PAM pattern.

### CRISPR Knock-in Designer pipeline design choice

The first stage in the CRISPR Knock-in Designer workflow is to perform codon mutations, which is done by mutating the target codon to all possible codons of the mutant AA or all Stop codons in case of nonsense mutations. The exact codon for the mutant AA residue may not matter to most users; if it does, the user can select a design with a specific mutant codon. In the next stage, we apply the CRISPR type option as well as all oligo options to select and run the correct design pipeline (Fig. 1). There are four such pipelines based on the choice of “Mutate PAM/sgRNA?” and “Silent mutations for restriction enzyme sites?” options. Cas9 designs can utilize all four of these pipelines, whereas Cas12a designs can run two of them that do not involve the PAM/sgRNA mutations procedure, as explained above (Fig. 1). Each run will utilize only one of the pipelines based on the options as described.

### Algorithms for additional mutations design

The best way to prevent re-cutting by a Cas9 enzyme after it mediated a genomic modification is to ensure that its guide RNA PAM site is mutated. For the protein-coding genes on which we are focused, another constraint is to ensure that the PAM mutation does not introduce further unintended missense mutations and does not interfere with exon splicing. Therefore, we have chosen synonymous mutations of codons overlapping the non-redundant part of a PAM as the procedure to inactivate the PAM of a gRNA used for point mutation knock-in design (Fig. 3). Briefly, PAM-overlapping codons are selected and verified to lie within an exon and not to be identical with the target codon. These codon(s) are then mutated synonymously and if the resulting mutant PAM sequence does not match its usual pattern, we record such a mutation as successful and add it to candidate oligos. Alternatively, we select the two codons closest to the PAM that also overlap the gRNA spacer sequence and, if available, mutate them synonymously to ensure that the sgRNA binding will be inactivated (Fig. 3). These mutation procedures are applied only to the Cas9 but not to the Cas12a designs (Fig. 1) due to the 18-nt distance between the PAM sequence and cut site of Cas12a enzymes. However, this distance also makes it very likely that the target codon overlaps the crRNA spacer, and its mutation will inactivate subsequent binding of Cas12a-crRNA complex and re-cutting.

**Figure 3.**
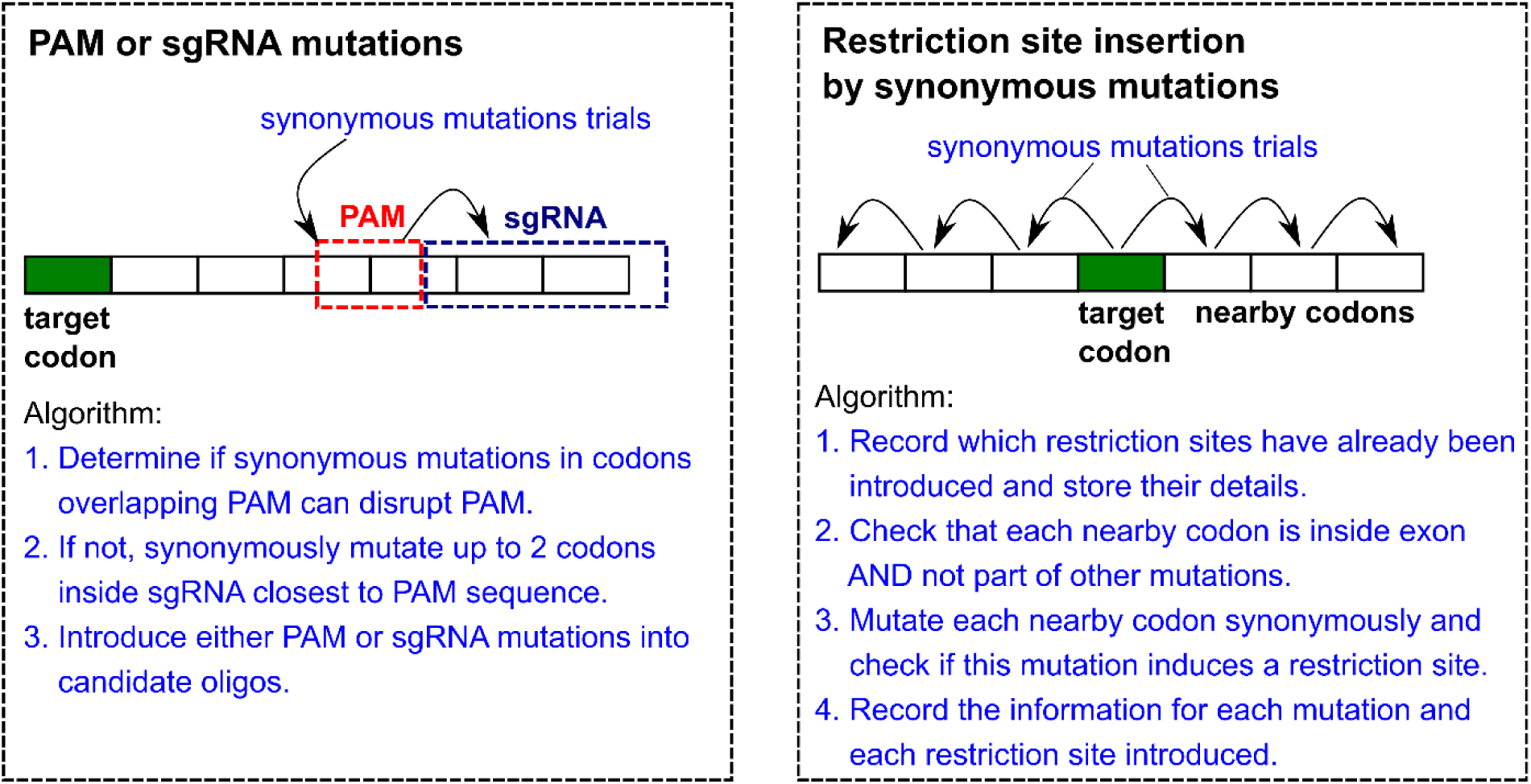
Algorithms for additional mutations selected in the oligo options. The algorithm for including PAM or sgRNA mutations is outlined where the focus is on the codons overlapping the PAM sequence and where synonymous mutations can inactivate it; an alternative is to inactivate the first two sgRNA spacer overlapping codons. Restriction site insertion by synonymous mutations is performed at one of the three codons on each side of the target codons depending on which of them are available. A detailed algorithm is described under the diagram. Target codon is marked in green, other codons are white rectangles, and the flow of mutation trials is indicated by the arrows.

Another useful feature of the CRISPR Knock-in Designer website is the ability to introduce restriction sites for genotyping purposes as well as to verify the introduction of sites by other mutations. First, all the pipelines verify if any other mutations have introduced restriction enzyme sites as their side effects and that any of these sites are unique in the genomic region spanned by the site assay primers. In case the user has selected the option to introduce restriction sites by synonymous mutations, the algorithm focuses on the maximum of 3 neighbouring codons on each side of the target codon (Fig. 3). Each of these codons is verified to lie within an exon and not to be part of other mutations. Synonymous mutations are introduced in each of these codons and then verified whether they introduce a unique restriction site. This is done by only focusing on the enzymes whose sites are absent from the site assay amplicon (non-cutter enzymes). Whenever a mutation generates a site/sites for one or several formerly non-cutter enzymes, both the mutation and enzyme information is recorded and stored in the resulting oligos. This algorithm has been inspired by the WatCut website (http://watcut.uwaterloo.ca/), but our implementation is unrelated and explicitly designed for this website.

### CRISPR Knock-in Designer output

The aim of any successful bioinformatics program is to produce correct and meaningful output. In the case of CRISPR Knock-in Designer, our goal has been to design oligonucleotides that can be used in experiments aimed at point mutation knock-in generation using CRISPR/Cas9 systems in protein-coding genes. The program output consists of two parts: TARGETING STRATEGY OUTLINE (Fig. 4A) and RESULTS OF THE OLIGO DESIGN (Fig. 4B) sections. The TARGETING STRATEGY OUTLINE section shows the location of site assay primers in the exon and flanking sequences, exon and intron sequences within the site assay amplicon, target codon, as well as the guide RNA and PAM locations, as shown for an example in Fig. 4A. Above the main section for design output is RESULTS OF THE OLIGO DESIGN which contains a variable number of oligo data output containers (Fig. 4B). Each of these containers contains the oligo sequence with target codon, PAM and mutations marked as in Fig. 4A. Restriction sites are also shown under the sequence as well as the sizes of fragments that these enzymes generate when digesting the site assay amplicons. Next, forward and reverse allele-specific PCR (AS-PCR) primers are shown in tables followed by the corresponding AS-PCR assays shown in a table below the primer tables (Fig. 4B). All these sequences and tables can be downloaded in a single text file by pressing the “Download oligo designs” button (Fig. 4B). The resulting text file contains identical information as the HTML output page except the mutated residue labeling as shown in an example (Fig. 4C). The user can then choose the most relevant oligonucleotide for them based on their preferences for the types of mutations and genotyping strategies. We also provide download buttons (“Download PrimeDesign inputs” and “Download pegFinder inputs”) for longer sequences prepared for PrimeDesign and pegFinder Prime Editing software tools (Fig. 4B). These sequences directly correspond to the designed oligos and have the same identifiers.

**Figure 4.**
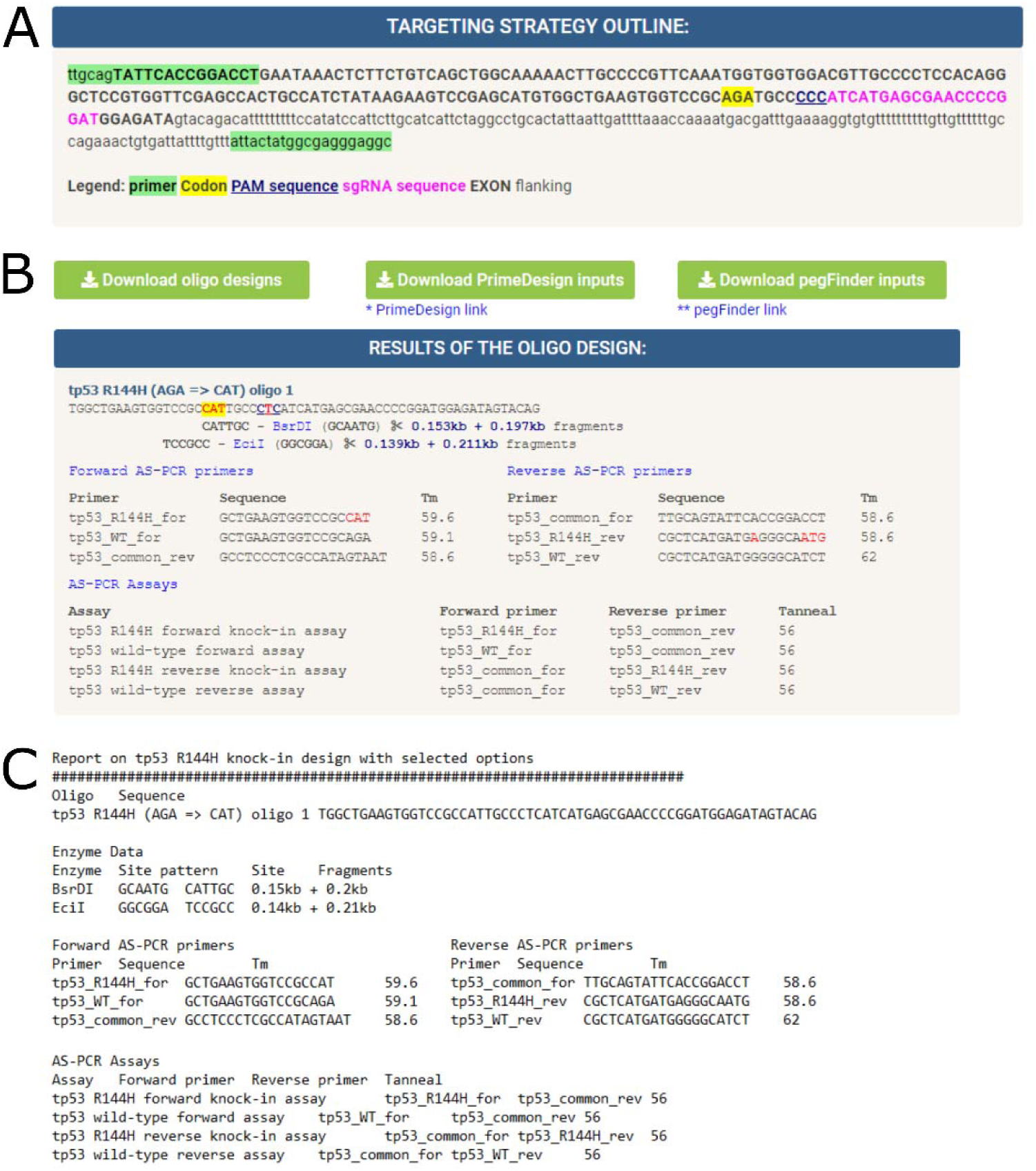
CRISPR Knock-in Designer outputs. The website generates two main types of outputs: targeting strategy (**A**) and oligonucleotide information (**B, C**). (**A**) TARGETING STRATEGY OUTLINE box shows a visualization of where all the relevant sequence elements are in the user provided sequence. These include primers used to select the relevant sequence region, sgRNA, its PAM sequence, target codon and whether a particular part of the sequence belongs to intron or exon. (**B**) RESULTS OF THE OLIGO DESIGN section typically contains file download buttons above the section header and multiple HTML containers for the designed oligos. The button “Download oligo designs” saves all the oligo designs in a single file. The other two buttons “Download PrimeDesign inputs” and “Download pegFinder inputs” enable downloads of longer input sequences suitable for the respective Prime Editing strategy designs containing the same mutations as the corresponding oligos. An example of an HTML container is shown here, and it contains the oligo name, sequence, the enzymes that can be used for genotyping, their resulting fragments as well as primers that can be used for forward and reverse AS-PCR assays. The correct primer combinations for these AS-PCR assays are shown at the bottom of the container. (**C**) The text version of the container shown in (**B**) from the file downloaded by pressing “Download oligo designs”.

### Software and code availability

The code for the main program and its several versions with pre-loaded data can be downloaded from the CRISPR Knock-in Designer GitHub repository (https://github.com/SergeyPry/knockinDesigner). GitHub README page describes how to run the website locally, either after the manual download or by automatically running the code from GitHub. The main web-based version of CRISPR Knock-in Designer on its own server with unlimited availability can be accessed at http://www.knockindesign.net/shiny/knockinDesigner/. This website version has its own GitHub repository: https://github.com/SergeyPry/shiny-server. The alternative website with limited active hours can be found at https://crisprtools.shinyapps.io/knockinDesigner/.

## Results

### Testing CRISPR Knock-in Designer supports its flexibility and robustness

To ensure that the final website functions without significant errors upon valid input, we have performed extensive testing. The initial testing was performed at the level of input data validation when for each input, a series of correct and incorrect inputs were generated and tested. Having fixed any code issues at this level, we have generated two locally run applications with embedded data (*tp53* R144H and *lmna* R471L zebrafish knock-ins) and the same server code to test new features, functions and workflows as the development progressed. The next series of tests was on 13 zebrafish point mutation knock-ins with multiple different guide RNAs (SupplementalFile1.txt), most of which we performed experimentally in the lab, in both manual and automatic modes. This testing resolved multiple issues related to the input guide RNA types and showed that the manual data input allows for a faster design upon submission, but the time needed to prepare and insert sequence data likely outweighs this benefit. Manual data input may also be beneficial if there is a large amount of variation in the targeting region such that the reference genome may not be fully applicable to the strain being used for the experiment. On the other hand, manual data input is essential for species not covered by the Ensembl REST API framework thus allowing for complete generalizability of the tool. We found that automatic retrieval from BioMart was rather slow during the initial testing, frequently resulting in >1 minute response time. Therefore, this approach was replaced with the Ensembl REST API-based retrieval, which has the same functionality but is at least 3-4 times faster when tested locally. Once the resulting issues were fixed, we expanded testing to 78 additional mutations in 50 genes from 5 species, most of which had multiple guide RNAs resulting in 179 testing cases (SupplementalFile2.xlsx). This set of testing cases includes two-exon straddling codons of different phases and from all 5 species to demonstrate the utility of the program for this challenging albeit rare type of point mutation knock-in design. These primary test cases were also evaluated in terms of the oligo output options, which are inclusion of additional mutations, orientation, and length of oligos. The current version of the website executed correctly on all the test cases in the current dataset. While optimizing the code, we encountered some errors for correctly performing gRNA and PAM mutations, exons including non-coding regions or single-exon genes, small exons (around 10 base pairs) mapping to multiple locations within messenger RNAs, retrieval of sequence data for large genes as well as some other less serious issues, all of which have been fixed by more robust code. Given that CRISPR Knock-in Designer is based on a deterministic algorithm and is not a machine learning application, we cannot provide a precise measurement of the algorithm accuracy, but the strategy overview output and oligo labeling allow the users to quickly discern any potential issues. The accuracy for future knock-in designs will be assessed empirically in our own usage as well as upon the user feedback and the code will be further improved if necessary.

### Prevalence and theoretical mutability of codons straddling two exons

To measure the prevalence of codons straddling two exons, we performed their direct quantification in protein-coding transcripts of six animal species (*Caenorhabditis elegans, Drosophila melanogaster, Ciona intestinalis, Danio rerio, Mus musculus*, and *Homo sapiens*) as a percentage of total coding sequence amino acids using exon phase data as well as transcript information. Two-exon straddling codon percentages were quantified by counting the phases for all exons of a transcript that are not equal to 0 and dividing this by 2 (types of phases), followed by the division with a total number of codons and multiplication by 100. The percentages were then averaged at the gene level by taking the mean of values for different transcripts of the same gene. We visualized the distribution of these percentages for all analyzed species (Fig. 5) as histograms where the x-axis is transformed by the square root and the y-axis represents the frequency of genes belonging to a particular group in the histogram. All species had from ∼3000 to ∼5000 genes without any two-exon straddling codons and the mean percentage of such codons was lower in invertebrates (0.68% in *C. elegans* and 0.35% in *D. melanogaster*) than in the chordates (vertebrates and the Urochordate *Ciona*) which all had a mean of ∼1% (Fig. 5). This analysis provides a snapshot of the prevalence of two-exon straddling codons and can be readily extended to any other genome where the data is available at both the aggregate and gene-focused levels.

**Figure 5.**
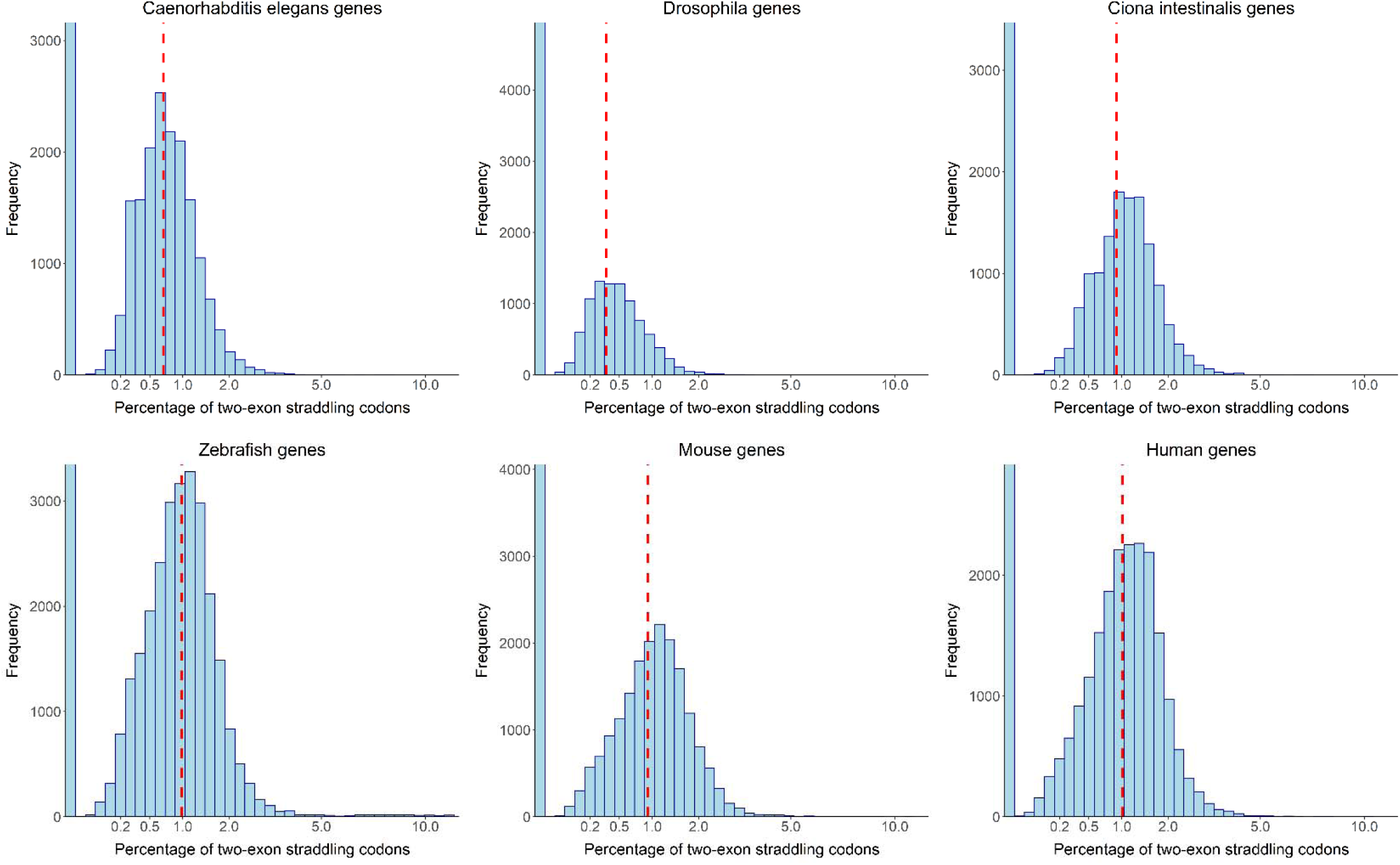
Quantification of the percentages of two-exon straddling codons in six animal genomes. Transcripts coding for proteins from six diverse animals (*Caenorhabditis elegans, Drosophila melanogaster* (Drosophila or fruit fly), *Ciona intestinalis* (a Urochordate species), *Danio rerio* (Zebrafish), *Mus musculus* (Mouse) and *Homo sapiens* (Human)) were assessed for the percentages of codons straddling two exons. These percentages were then averaged at the gene level and visualized as histograms. To provide more detail of the distribution, the horizonal axis was scaled by the squared root of the values. The relevant percentage values are indicated on the horizontal axis and the mean value for each distribution is shown by the dashed red line.

Although the mean prevalence of codons overlapping two exons is relatively low at 0.35-1 % (Fig. 5), they may encode important amino acid residues. CRISPR Knock-in Designer is designed to handle situations where a codon overlaps two exons. To mutate such codons, a mutation needs to be introduced into either of the relevant exons. Therefore, the gRNA site strongly influences the possibility of mutating codons overlapping two exons because the cut site determines which exon can be mutated. To address the question of codon mutability and facilitate the computational design process, we have introduced codon fragment phases, which are +1 and -2 for an intron phase 1 or +2 and -1 for an intron phase 2 (Fig. 6A). As a next step, we tested the mutability of each codon fragment in isolation and the combined (summed up) +1/-2 or +2/-1 codon mutability (Fig. 6B). As expected, +2 codon fragment had the highest overall mutability, followed by the -2 fragment and then by +1 and -1 codon fragments (Fig. 6B). Overall, +2/-1 codons were 65.2 % mutable suggesting they are more mutable than the +1/-2 codons (36.1 %) (Fig. 6B). These results illustrate the structure of the genetic code from the perspective of all possible mutations of codon fragments. However, the naturally occurring mutations in such codons will most likely arise by mutations in individual codon fragments rather than both fragments being mutated simultaneously. Thus, most of the natural mutations of two-exon straddling codons will correspond to the mutable (blue) boxes of the theoretical mutability plots (Fig. 6). However, some mutations needed for experimental purposes may not be mutable by local ssODN-based strategies and will instead have to be mutated by two-site knock-in strategies or larger-scale genome engineering approaches. For a particular two-exon straddling codon, CRISPR Knock-in Designer either outputs the designed oligos or a statement that the specific knock-in strategy is not feasible.

**Figure 6.**
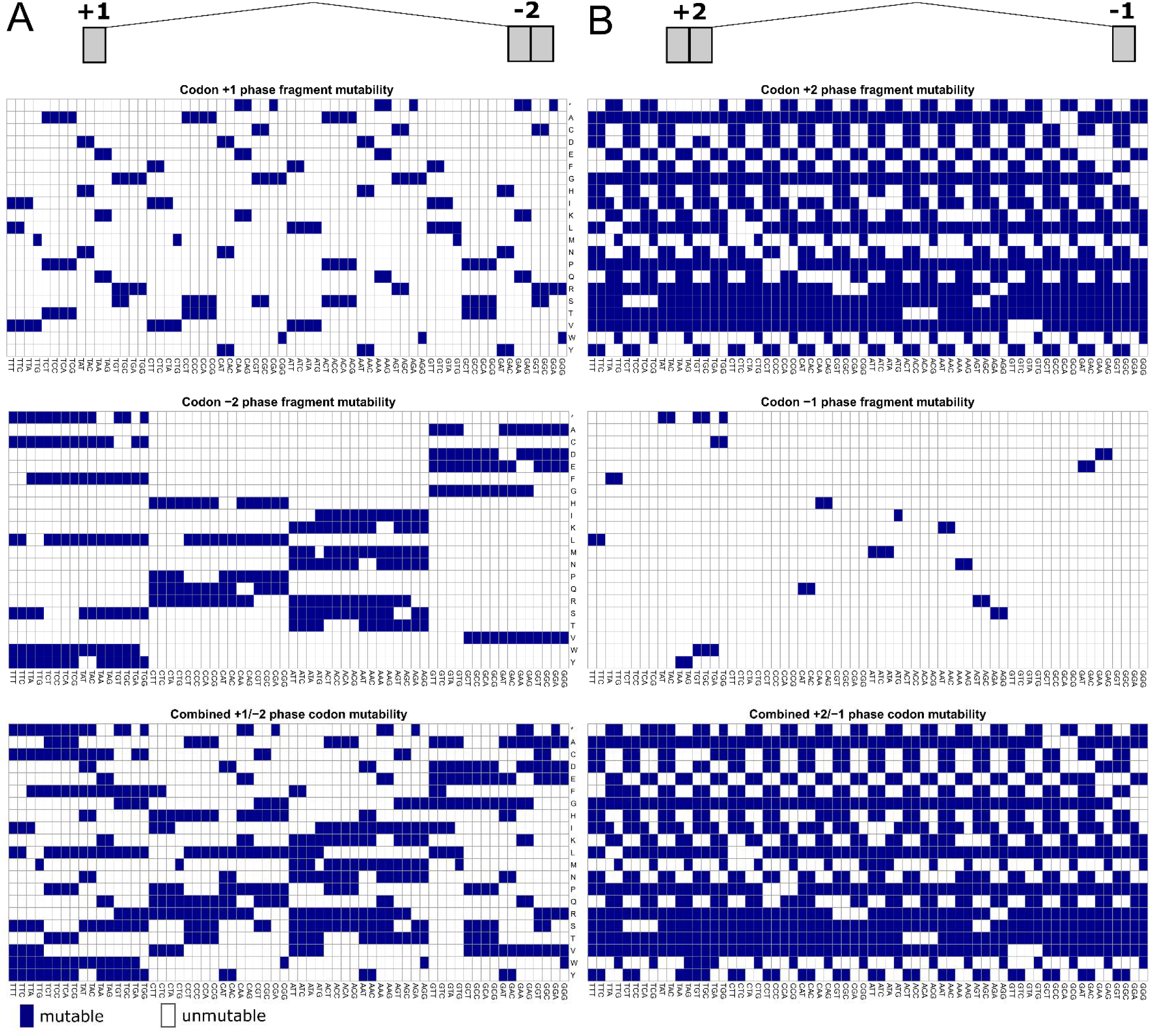
Mutability of codons straddling two exons to different amino acids. Codons straddling two exons cannot always be mutated to another codon encoding a particular amino acid. The +1/-2 codons (**A**) and +2/-1 codons (**B**) are shown as their nucleotides are arranged in the genome and their differences with respect to the mutability have been visualized as matrices of codons by possible amino acids (including stop codons indicated by ‘*’). (**A**) Mutability matrices for +1 phase and -2 phase codon fragments as well as the combined +1/-2 codon mutability (when the optimal fragment is chosen for mutation) are shown. (**B**) The same visualization is performed for +2/-1 codons and their fragments. Overall, these mutability heatmaps illustrate the mutability testing code implemented in CRISPR Knock-in Designer.

### Comparison to other point mutation knock-in tools

Currently, there exist two prominent ssODN donor knock-in design tools, IDT (https://www.idtdna.com/pages/tools/alt-r-crispr-hdr-design-tool) and Benchling HDR design tools (https://www.benchling.com/). Both the algorithms are commercial and require a login for access. Unlike our tool, both the above tools also offer a graphical user interface and can build gRNA within the platform. In contrast, our algorithm provides a minimalistic user interface and is quite simple to handle. All three tools offer support for both manual and ID-based user inputs.

CRISPR Knock-in Designer also offers additional features like restriction site introduction by synonymous codon mutation and primer design to carry out AS-PCR assays that neither of the other available tools provides. Uniquely, Benchling provides users with codon adaptation index calculation allowing the users to select codons very similar to the existing codons. CRISPR Knock-in Designer also offers an option to download the results in a user-friendly way, a feature absent in other tools. Moreover, we provide sequences to perform designs of the specific mutations for the prime editing approach, a new feature, not available in other knock-in design tools.

### pegRNA design based on the CRISPR Knock-in Designer output

Prime editing technology is based on a *Streptococcus pyogenes* Cas9 nickase enzyme fused to a reverse transcriptase that recognizes NGG PAM sequence (20). The currently available websites for pegRNA design rely on the user precisely specifying the nucleotide sequences to be modified. We sought to adapt the output of our program to serve as the input for either PrimeDesign (21) or pegFinder (22). Input for pegRNA design does not require PAM or gRNA spacer mutations because pegRNA design is independent of the gRNA input to CRISPR Knock-in Designer. Therefore, we recommend that “Synonymous codon mutations of PAM or gRNA spacer?” option have the value “No”, but the downloads for Prime Editing inputs will be provided for any design. These pegRNA design inputs will be provided to the user as files available from download buttons (“Download PrimeDesign inputs” and “Download pegFinder inputs”) at the top of the output (Fig. 4B). PrimeDesign input requires special formatting for nucleotide substitutions, whereas pegFinder requires a wild-type and a modified sequence to be provided separately. These files are FASTA files with the same names as the oligos to maintain the correspondence. An example pegRNA strategy design from a point mutation definition via the CRISPR Knock-in Designer workflow is shown in Figure 7.

**Figure 7.**
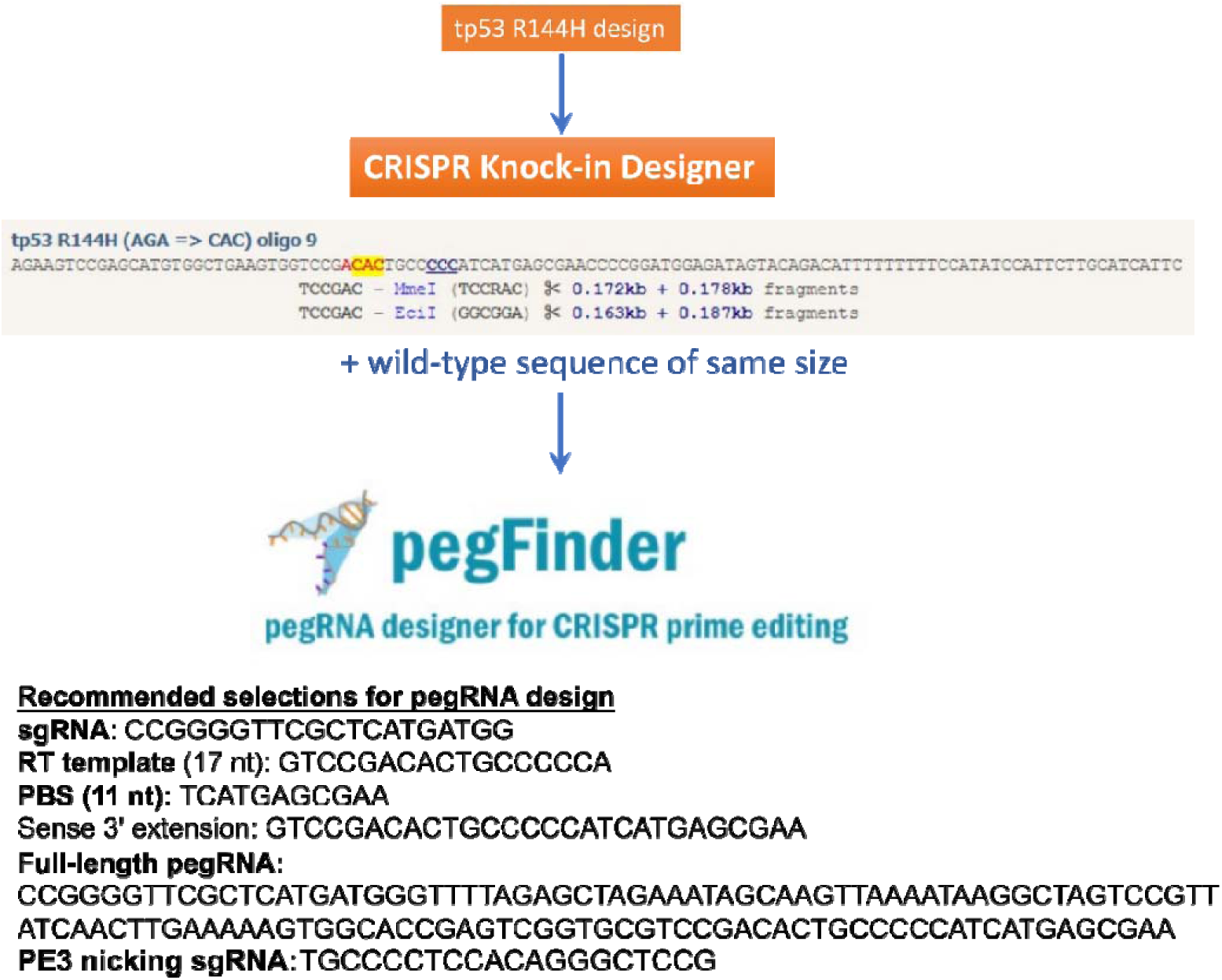
Design of pegRNA using pegFinder from the CRISPR Knock-in Designer output. Upon designing oligos using CRISPR Knock-in Designer, the user can download a file with the desired mutated and wild-type sequences around the site of mutation and use it to run pegFinder. pegFinder, in turn, provides the optimal parameters of its design and several alternative ones as well as additional “nicking” sgRNA for either PE3 or PE3b strategies.

## Discussion

The methodological advances in the techniques to introduce precise point mutations into protein-coding and other genes have mainly focused on the experimental optimizations for better efficiency and precision, more broad applicability to a wider range of species and the search for novel technologies to circumvent the challenges of performing point mutation knock-in using the standard CRISPR/Cas9. All these directions have resulted in significant successes. Most model species have protocols and optimization studies available to guide researchers in developing their strategies to introduce point mutations using CRISPR/Cas9 (8, 9, 12–14, 23–26). The weakness of these studies is that the design process for oligonucleotides is mostly manual with some aid of basic bioinformatics and other software tools. The currently available commercial knock-in oligonucleotide design tools do not go much beyond showing the relevant nucleotide and codon numbers, the same functionality also available in the DNA sequence editors. Genotyping strategies also must be designed manually in a series of potentially error-prone steps. With the CRISPR Knock-in Designer, we have sought to overcome these limitations by designing a “one-stop-shop” oligonucleotide design pipeline that minimizes the required data input and outputs, encompassing both the oligonucleotide sequences and a comprehensive set of genotyping strategies. The flexible options of our tool allow the users to further refine the parameters of the resulting oligonucleotides as described in this study. The search for novel technologies to introduce point mutations without the double-stranded breaks intrinsic to the CRISPR/Cas9 has also been fruitful. Base editors are nuclease-deficient Cas9 fusions with deaminase enzymes targeting cytidine, adenine, and other bases (27). Huge efforts to improve base editors have resulted in a large toolkit of efficient and precise enzymes for point mutations, which nevertheless have the limitation that they do not introduce multiple different mutations when one base editor is applied (28). By contrast, Prime Editing does not have strong limitations on the exact sequence of introduced modification, making it analogous to ssODN-based point mutation CRISPR/Cas9 knock-ins. We have therefore provided the option for the users to generate inputs for pegRNA design software alongside their oligonucleotide designs. The purpose here is to provide more options for generating the desired mutations by either CRISPR/Cas9-based knock-ins or, if these are inefficient, by prime editing. Given that the intended sequence replacement would be identical in both cases, the genotyping strategies designed will be valid for both technologies. Overall, it is likely that in the foreseeable future, CRISPR/Cas9-based knock-ins, base editors and prime editing technologies for point mutations will co-exist, suggesting that the software integrating the designs for these technologies will have great utility.

